# Loop-Directed Cofactor Binding Enables Programmable Multicofactor Protein Design

**DOI:** 10.64898/2026.06.27.734619

**Authors:** Andrew C. Mutter, Aleksandr Uvaydov, Eskil M.E. Andersen, Sara Morsi, Sarah Beck, Mohammad Khan, Bruce A. Palfey, Carolyn E. Lubner, Ronald L. Koder

**Author notes:** Merck, Inc., CBS-AR&D Biologics Potency Group., 126 E. Lincoln Ave., Rahway, NJ 07065. to whom correspondence should be addressed: Department of Physics, CDI 1.315, The City College of New York, 85 Saint Nicholas Terrace, New York, NY 10031. These authors contributed equally (Andrew C. Mutter, Aleksandr Uvaydov).

## Abstract

The emergence of respiratory, photosynthetic, and assimilatory complexes in evolution required proteins capable of binding multiple catalytic and electron-transfer cofactors while exerting fine control over their spatial arrangement. Across natural systems these cofactors are preferentially positioned in loop regions. In contrast, most protein design strategies have focused on installing cofactor-binding sites within helical elements. Here we show that introducing only two histidine ligands into the interhelical loops of a single-chain four-helix bundle creates an additional site that binds ferric heme with an equilibrium dissociation constant of 90 nM. This nanomolar affinity is achieved through a minimal sequence change, enabling assembly of complexes containing up to three distinct cofactors with positional specificity within one designed protein domain. Using this strategy, we created constructs containing one or two hemes in combination with Zn(II)–phthalocyanine monosulfonate, Zn-heme, and the light-harvesting Zn(II)–tetraphenylporphyrin tetrasulfonate. Fluorescence measurements of constructs containing the latter show efficient energy transfer between photoactive donor cofactors. By demonstrating that loop-embedded ligands support robust, modular, and evolutionarily plausible cofactor recruitment, this work provides a mechanistic explanation for the widespread placement of redox and catalytic cofactors in loops in natural proteins: only limited packing complementarity is needed, meaning that just a few mutations can introduce a functional cofactor binding site, after which additional mutations can tune affinity, reactivity, and specificity. More importantly, it establishes a straightforward path toward constructing functional protein domains that mirror the complexity of biological energy-conversion architectures.

## Introduction

The function of complex natural enzymes such as the cytochromes *bc_1_* and *b_6_f* or the photosystems is, to a first order approximation, a result of the protein scaffold holding multiple small molecule cofactors such as hemes, quinones, chlorophylls and carotenoids at the appropriate spacing to enable electrons to travel from one end, at which one active site extracts electrons from a substrate, to the other, at which a second active site uses that electron to perform reductive chemistry (1–4). These cofactor arrays are termed electron transfer chains. In natural photosynthetic proteins, light-activated charge separation begins at a primary donor cofactor located at the center of such an electron transfer chain, and the chains are organized such that the resultant electron and hole move away from each other toward the sites of their chemical action (5).

A number of groups, including ours, have been working toward the design of self-assembling cofactor-containing proteins as both a means of learning more about their evolved function (the maquette approach (6–9)) and a pathway toward useful functional catalysts (10–14). Our group has used bioinformatics to examine and create single natural heme (15; 16) and primary donor (17; 18) binding sites in designed proteins and has used these sites to create functional models of oxygen transport proteins (19; 20) and solar energy conversion proteins. Each of these binding sites were designed to bind cofactors via ligand residues which are located on alpha helices. Natural proteins, however, primarily bind cofactors in loop regions (21). To our knowledge, no designed heme- or porphyrin-binding endeavor has targeted cofactor binding to loops.

Single-site catalytic designs are only a starting point toward creating realistic models of complex multicofactor systems such as the photosystems, respiratory complexes, and assimilatory electron-transport chains. Several groups have built dyad proteins that support light-driven electron transfer (22–25) but proteins engineered to stably bind three or more redox-active cofactors remain uncommon. McAllister and colleagues (26) created a helical bundle that binds four identical Fe(II)–porphyrins and Cohen-Ofri et al. re-engineered HP7 to accommodate three identical Zn(II)–bacteriochlorophyllide cofactors at two sites (27). A major advance came from Ennist and coworkers, who assembled three chemically distinct cofactors—a heme, a photoactive Zn-porphyrin, and a di-Zn or di-Mn cluster— within a single extended helical scaffold (28). Here we extend this trajectory by modifying a single-chain, two-cofactor tetrahelical bundle to position a third cofactor-binding site within the loops connecting helices 1–2 and 3–4. This simple architectural change supports assembly of several distinct heterocofactor combinations, enabling the simple construction of more sophisticated multifunctional constructs than previously possible.

## Results and Discussion

### Protein design

The two-cofactor-binding, single-chain, four-alpha-helix-bundle protein *HHHF* was chosen as a starting point because it is stable, soluble, well characterized, and already binds two different porphyrin-like cofactors with positional specificity: one iron-containing cofactor which binds via hexacoordinate bis-histidine ligation at the open end of the helical bundle, and one zinc-containing cofactor which binds using pentacoordinate mono-histidine ligation near the end of the protein which contains two connecting interhelical loops (Figure 1B) (18). The distance between the cofactors has been modeled to be 17 Å. Each helix in this protein has a tryptophan three residues N-terminal to the ligand histidine (see Figure 1), a position which is expected to bring the side chain into contact with the porphyrin ring of the heme cofactor (29). Tryptophan side chains have been shown to quench fluorescence in proteins both by energy-transfer and electron-transfer mechanisms,(30) and for this reason we changed the two tryptophans predicted to pack against the zinc-porphyrin to photo- and redox-inactive phenylalanine residues. *Photo* binds a single ferrous heme cofactor 3-fold weaker than *HHHF* (Figure 2A), and a bound heme reduction potential 50 ± 20 mV lower (Table 1 and Figure 2C). The ferric heme affinities of both proteins are also within error of each other.

**Figure 1.**
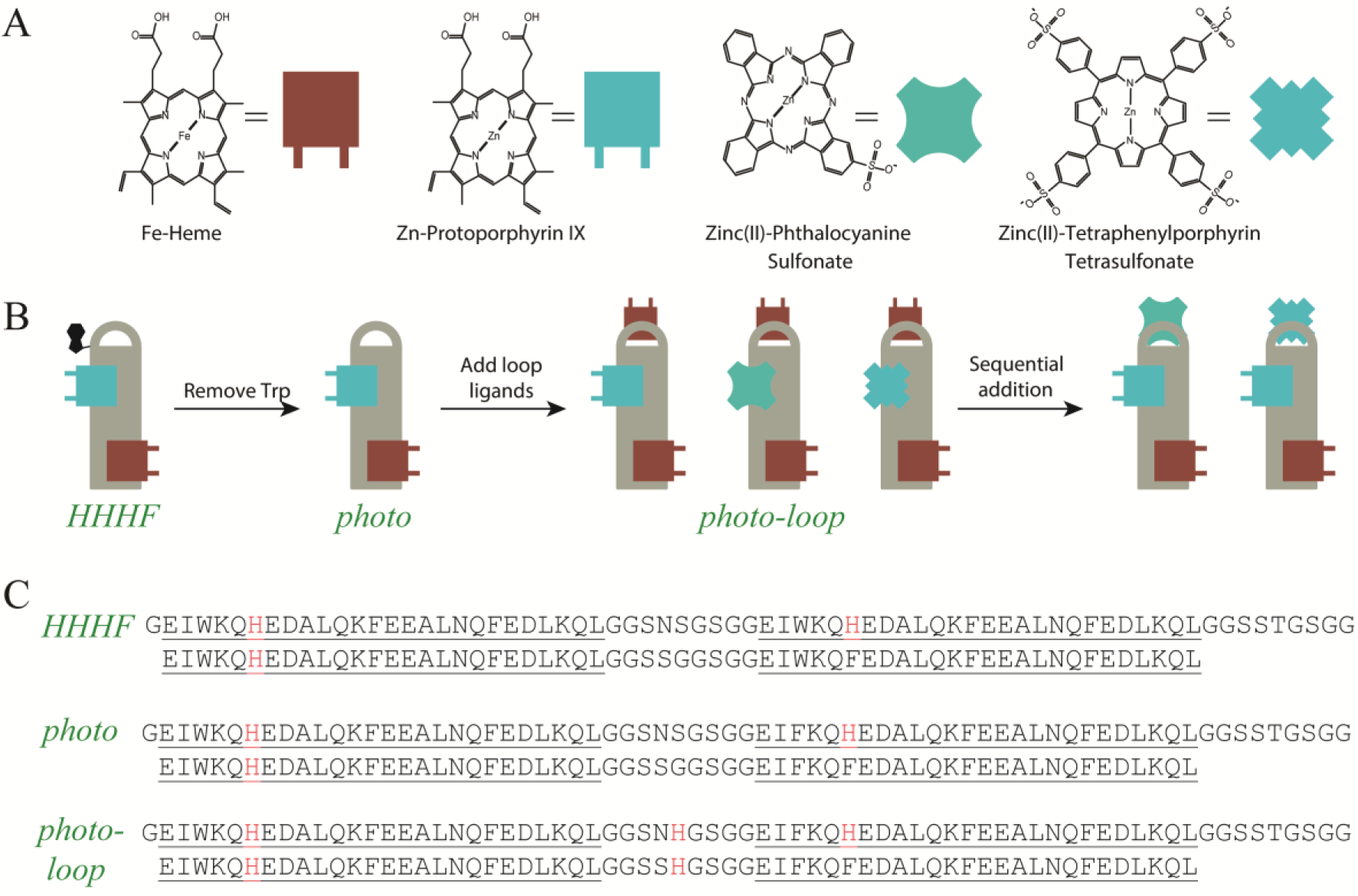
Schematic overview of the design pathway used to make heterotricofactor proteins. (A) Structure of the cofactors used. (B) The design pathway leading to the photo-loop protein. (C) Primary structure of each protein used in this work.

**Table 1.** Thermodynamic and kinetic parameters for bicofactor complexes.

| Protein | K<sub>d,red_heme</sub><br>(nM) | E<sub>m</sub> <sup>#</sup><br>(mV) | K<sub>d,ox_heme</sub><sup>\$</sup><br>(nM) | k<sub>on,ox_heme</sub><br>(mM<sup>-1</sup>s<sup>-1</sup>) | k<sub>off,ox_heme</sub><sup>↓</sup><br>(s<sup>-1</sup>) | K<sub>d, Zn-heme</sub><br>(nM) | Fluorescence<br>(rel intensity) | k<sub>on Zn-heme</sub><br>(mM<sup>-1</sup>s<sup>-1</sup>) | K<sub>off Zn-heme</sub><br>(s<sup>-1</sup>) |
|---|---|---|---|---|---|---|---|---|---|
| *HHHF* | 220±20<sup>*</sup> | -260±9<sup>*</sup> | 0.20±0.02 | 230±20 | (4.6±0.5)x10<sup>-5</sup> | 90±20 | 1.0 | 9±2 | (7.2±0.7)x10<sup>-3</sup> |
| *photo* | 700±200 | -311±8 | 0.18±0.05 | 380±20 | (6.8±0.7)x10<sup>-5</sup> | 300±90 | 1.46 | 3.8±0.5 | (4.8±0.4)x10<sup>-2</sup> |

<sup>\*</sup> Data taken from reference 19.
<sup>#</sup> Midpoint reduction potential.
<sup>\$</sup> Dissociation constant in the ferric state calculated using thermodynamic cycles, see reference 8.
<sup>↓</sup> Off-rate calculated using K<sub>d</sub> and k<sub>on</sub>

**Figure 2.**
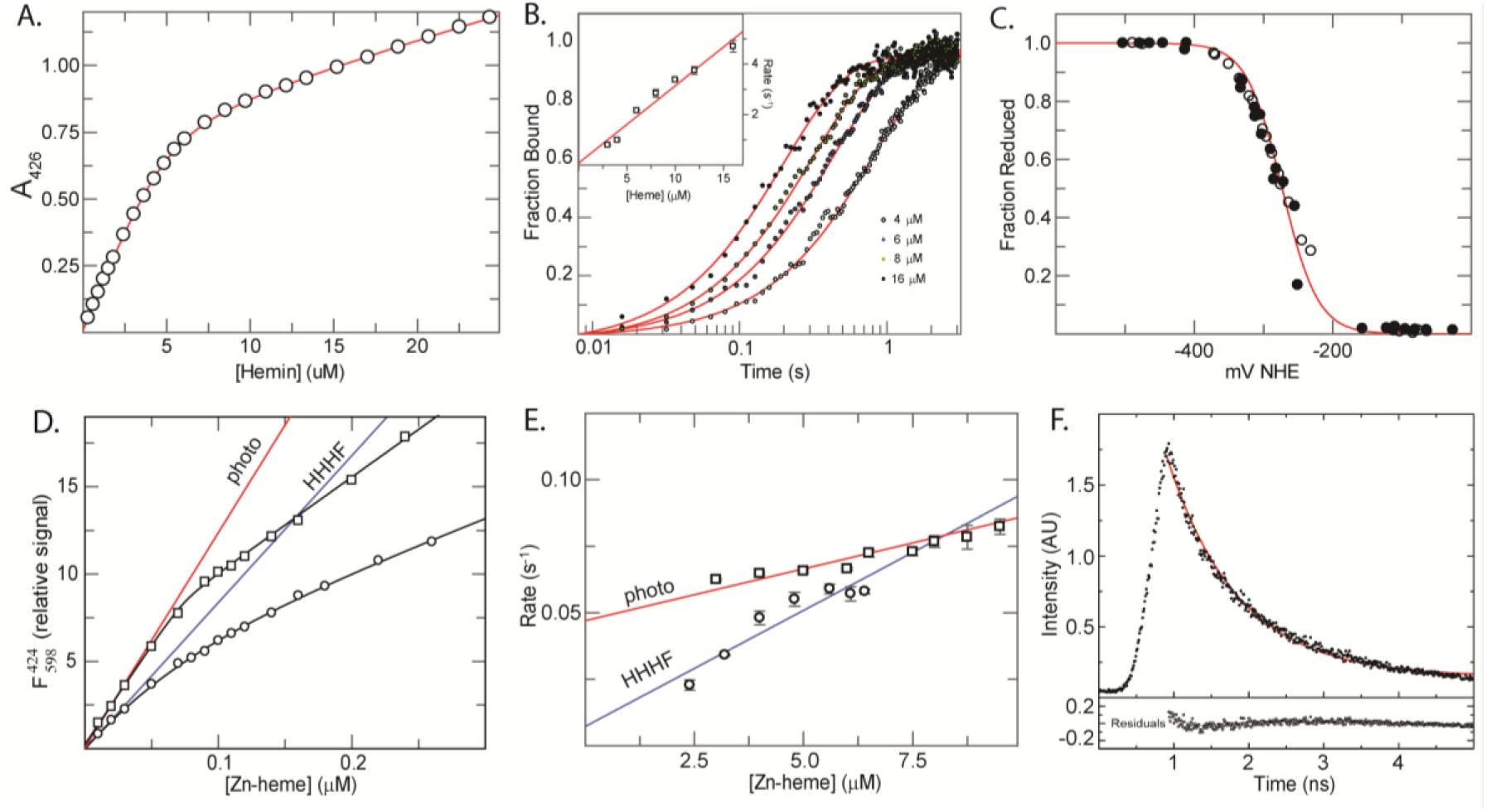
Heme and primary donor binding to *HHHF* and *photo*. (A) Optical binding titration of heme to *photo* under reducing conditions. Points are the absorbance at the reduced Soret maximum *vs.* the number of equivalents of heme. Line drawn is a fit with equation 3 using a Kd of 700 nM. The break is at 0.97 equivalents. (B) Selected time courses of heme binding to *photo*. Inset is a replot of binding rates as a function of heme concentration. Line drawn is a fit with equation 5 with a kon of 380 mM^-1^s^-1^. The intercept, or koff, goes through the origin within error. (C) Potentiometric titration of the *photo*:heme complex. Filled circles are points taken in the reducing direction, open circles are points taken in the oxidizing direction. The line is a best fit to all of the data points with equation 7 using a single reduction potential of -310 mV *vs.* NHE. (D) Fluorescence titration of the *HHHF*:heme and *photo*:heme complexes with Zn-PPIX. Curving lines are a best fit of the emission intensity at the λmax with equation 4 with dissociation constants of 90 and 200 nM respectively. Straight lines are a linear fit of the first 0.2 equivalents with slopes of 76 and 111 μM^-1^, respectively. (E) Replots of the binding rates *vs* Zn-PPIX concentration for both the *HHHF*:heme and *photo*:heme complexes. Error bars are standard deviations of three separate binding experiments. Lines drawn are linear fits with equation 5 using kon values of 9 and 3.8 mM^-1^s^-1^ and koff values of 0.0072 and 0.048 s^-1^ respectively. (F) Fluorescence lifetime of the *photo*:heme:Zn-PPIX bicofactor complex. Red line is a single exponential fit of the decay curve with a lifetime of 1.06 ns. Lower line is the residual of the fit.

To better analyze cofactor binding, we used pre-steady-state kinetics to determine heme binding rates to the bis-histidine site of each protein. The ferric heme association rate is 65% faster to *photo* than to *HHHF* (Figure 2B), likely a result of the lowered stability and increased dynamic motion of the former (31). For both proteins the dissociation rate is too small to be directly determined from on-rate measurements (see eqn 3), but can be derived using measured on-rates in combination with binding affinities: the observed off-rate constant is 48% faster from *photo* than from *HHHF* (Table 1).

We next quantified Zn-heme binding to the pentacoordinate sites of both *photo* and *HHHF* and compared fluorescence and excited-state electron transfer behavior of the bound cofactor. Figure 2E depicts the concentration-dependent binding rate of Zn-heme to *photo* and *HHHF*. In the case of heme acceptors, the ratio of the fluorescence quantum yields observed when the heme is in the ferric and ferrous states is commonly used to measure the fractional electron transfer yield (18). When tryptophan is the acceptor, however, it is not possible to make a similar comparison, as the tryptophan reduction potential is outside the range at which it is possible to chemically affix the water solution potential (32). Instead, we have performed Zn-PPIX binding titrations both to heme-containing *HHHF* and to *photo* and used the initial slopes in the binding titrations to determine their fluorescence quantum yields (Figure 2D). Zn-PPIX binds almost two-fold more weakly to the pentacoordinate binding site of the ferrous heme-*photo* complex, while the binding rate is two-fold slower. Zn-PPIX bound to *photo* is 46 ± 5% more fluorescent than when bound to *HHHF*. The increased fluorescence is consistent with the removal of the indole electron acceptor: the ratio of bound fluorescent quantum yields indicates that almost one-third of the absorbed photons in *HHHF* generate electron transfers to the tryptophan indole side chain, and *photo-*ZnPPIX complexes represent a significant improvement in terms of quantum yield.

To more accurately quantify the electron transfer and energy transfer properties of *photo* and its complexes, we measured the fluorescence lifetime of the *photo-*heme-ZnPPIX complex using time-resolved photoluminescence spectroscopy. As figure 2F depicts, in the absence of an acceptor tryptophan or cofactor the fluorescence lifetime is 1.06 ± 0.01 ns. This lifetime in combination with the relative steady-state intensities from 2D enables us to use Eqn. 8 to calculate the electron transfer rate to tryptophan in *HHHF*: (4.3 ± 0.2) x 10^8^ s^-1^.

### Addition of a third binding site

The ferric heme-histidine interaction energy has been estimated to be 6 kcal/mol for each histidine in a bis-histidine ligation site (33). 12 kcal/mol is sufficient energy to bind a heme even in the absence of additional stabilizing interactions. This led us to create a very simple third cofactor binding site by adding two histidine residues to the glycine-rich loops that we used to connect helices 1 to 2 and 3 to 4 (Figure 1C), creating a protein that we have termed *photo-loop*.

Photo-loop derives from a binary-patterned scaffold developed before the advent of AlphaFold and was neither selected nor optimized using structure-prediction confidence scores. We used AlphaFold3 (34) retrospectively to examine possible cofactor arrangements within this scaffold. Models containing three heme molecules placed the cofactors at the three intended sites: the original bis-histidine heme site, the internal mono-histidine porphyrin site, and the newly introduced loop-histidine site (Figure S1). The buried cofactors had higher mean pLDDT values (58 ± 1) than the loop-bound heme (47 ± 2). These confidence values should be interpreted in the context of a sequence-divergent de novo protein whose design was independent of AlphaFold, which have been found to often be significantly lower than those designed using it (35). The models provide a plausible structural arrangement consistent with the intended architecture, with greater uncertainty in the precise placement of the loop-bound cofactor. The binding measurements below provide the experimental evidence for recruitment of an additional heme.

Kinetic analysis of ferric heme binding to the loop binding site gives a k_on_ of 210 ± 40 mM^-1^s^-1^ and a k_off_ of 0.9 ± 0.5 s^-1^ (Figure 3C). The ratio of these rates gives a dissociation constant of 4 ± 2 µM. As we have shown before (16), heme binding can be a two-step process, where the initial singly ligated pentacoordinate complex observed in stopped-flow is followed by a unimolecular rearrangement in which a second, distal histidine ligates the heme iron (Figure 3E) and the equilibrium binding constant is:

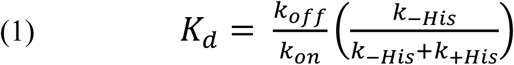

**Figure 3.**
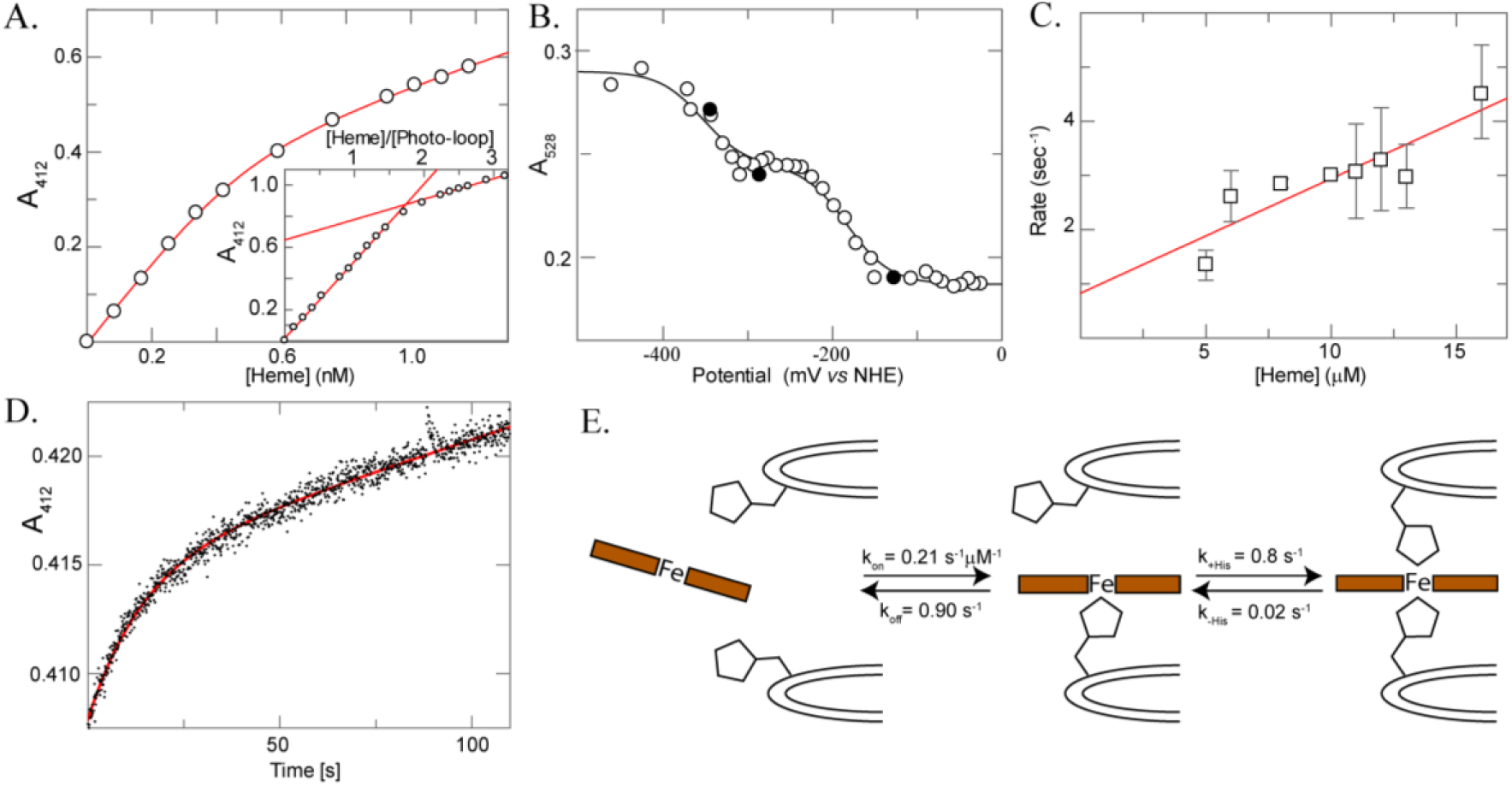
Heme binding to *photo-loop*. (A) Equilibrium binding titration of the second heme to 500 nM *photo-loop* under oxidizing conditions. Points are the absorbance at the ferric Soret maximum *vs.* the number of equivalents of heme. Line drawn is a fit with equation 3 using a Kd of 90 nM. *Inset.* Full oxidized binding titration under the same conditions. The break is at 1.8 equivalents. (B) Potentiometric titration of the *photo-loop*:diheme complex. Filled circles are points taken in the reducing direction, open circles are points taken in the oxidizing direction. The line is a best fit to all of the data points with equation 7 using two reduction potentials of -350 mV and -185 mV vs. NHE. (C) Rates of binding of heme to the loop binding site in the heme:*photo-loop* complex. Line drawn is a fit with equation 5 with a kon of 0.210 µM^-1^s^-1^ and an intercept, or koff, of 0.9 s^-1^. (D) Heme dissociation from the loop binding site of diheme *photo-loop.* Line drawn is fit with equation 6 with an apparent rate constant, koff,app of 0.071 s^-1^. (E) Schematic of the two-step binding of heme to the loop binding site with experimental and derived rates for each step.

Oxidized hemes do not bind carbon monoxide, so laser flash photolysis techniques will not serve to extract distal histidine association and dissociation rates. We instead measured the apparent heme off-rates, k_off,app_ by mixing diheme *photo-loop* with apomyoglobin (36).

Diheme *Photo-loop* releases one heme with a rate *k_off,app_* of 0.071+0.002 s^-1^ (Figure 3D), and one with a rate on a tens of minutes timescale. Off-rates are the same within error when the experiment is performed using 26 or 52 µM apomyoglobin, indicating that in these experiments heme binding to apomyoglobin is not rate limiting. Control experiments using heme:*photo* complexes demonstrate that the slow off-rate is the helical heme binding site. Ligand dissociation in two-step binding processes such as this is given by (37):

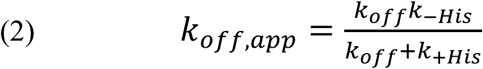

which in combination with equation 1 enables the calculation of k_+His_ and k_-His_ rates of 0.8 ± 0.5 s^-1^ and 0.02 ± 0.01 s^-1^ respectively (see derivation in Figure S2). These values predict an equilibrium dissociation constant of approximately 0.10 µM, consistent with the measured value of 0.090 ± 0.030 µM. These parameters predict an apparent heme dissociation rate of approximately 0.011 s⁻¹, within an order of magnitude of the independently measured rate of 0.071 ± 0.002 s⁻¹. This agreement across equilibrium and kinetic measurements is consistent with a mechanism in which initial heme binding is stabilized by subsequent distal-histidine ligation. The ratio of *k*_+His_/*k*_−His_ ≈ 40, indicating strong stabilization by distal histidine ligation.

### Creating photoactive ternary complexes

As Figure 3A demonstrates, *photo-loop* binds two heme cofactors at the two flanking bis-histidine coordinating sites, leaving the central monohistidine site open to bind a pentacoordinate metalloporphyrin ligand. Starting with the diheme *photo-loop* complex, we created three diheme heterotricomplexes by binding ZnPPIX, ZnPcS and ZnTPPS_4_ at this central pentacoordinate binding site (Figure 4 and Table 2)). All three of these Zn-containing porphyrin cofactors are capable of performing as light-activated electron-donating molecules. We have previously created *HHHF* heterocomplexes containing one heme and either a single ZnPPIX or ZnPcS and have determined that there is no light-activated electron transfer between the cofactors, likely because of the large distance between the two binding sites (18). As Figure 4 demonstrates, the proximate location of the loop binding site brings the second heme cofactor much closer, resulting in significant quenching of the Zn-cofactors. The ratio of the ferric to ferrous Zn-cofactor fluorescence demonstrates 58%, 5% and 29% quenching in the case of Zn-PPIX, ZnPcS, and Zn-TPPS_4_ cofactors respectively. A similar result is seen when Zn-PPIX is titrated into diheme *photo-loop* (Figure S3). However, at distances as short as this, spin-orbit relaxation is an alternative quenching mechanism (38), and given the relative reduction potentials (39) and the single-digit nanosecond fluorescence lifetimes of the bound Zn cofactors, it is possible such relaxation plays some part in the observed loss of fluorescence. However, we can use Equation 5 to combine these quenching values and the fluorescence lifetimes of the bound Zn cofactors in the absence of the second heme to predict upper limits for the electron transfer rates of each heterocomplex: 1.3×10^9^, 2.7×10^8^and 5×10^7^ s^-1^ for Zn-PPIX, ZnTPPS_4_, and ZnPcS respectively (Table 2).

**Figure 4.**
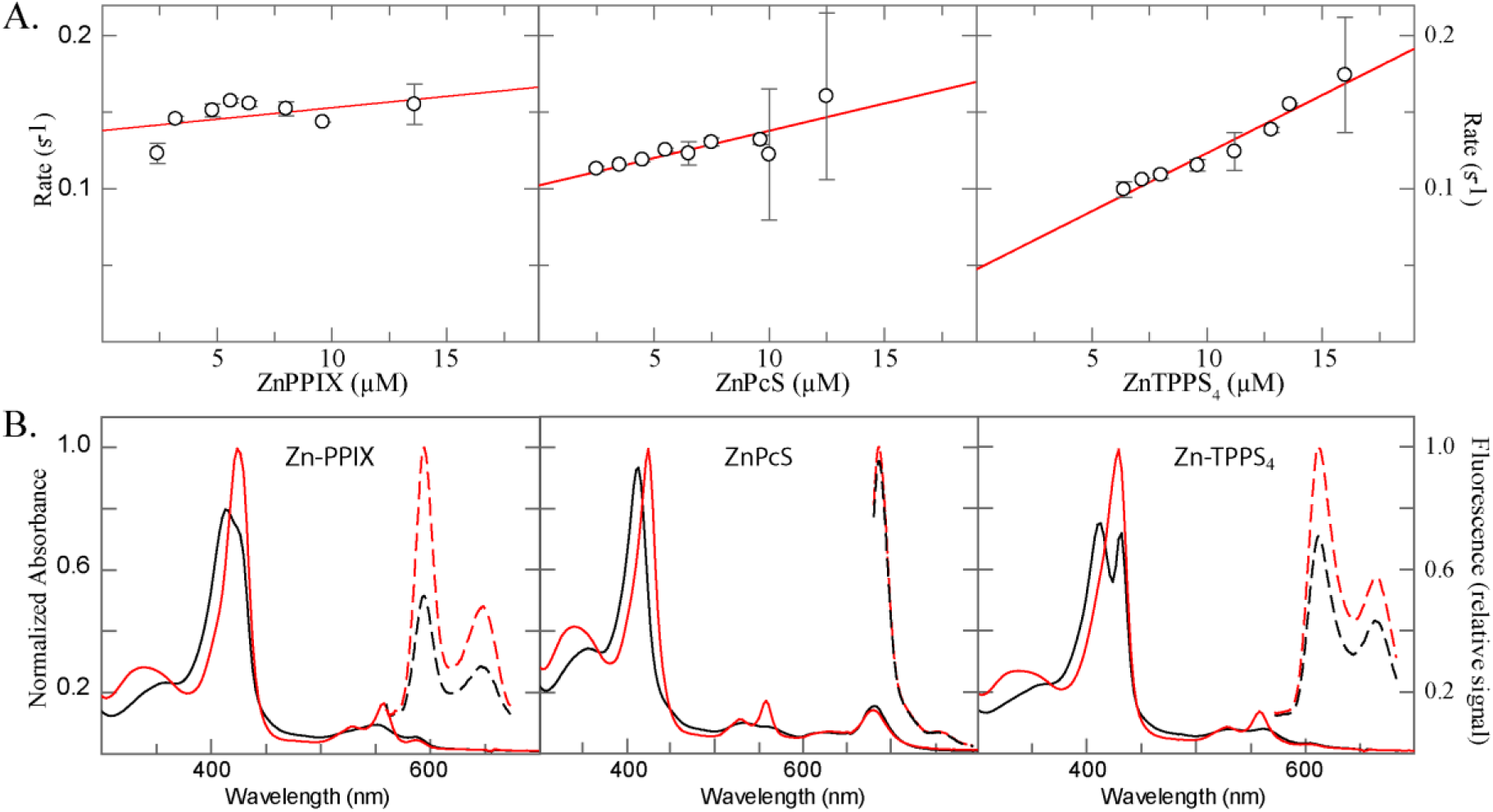
Zn-cofactor binding to diheme *photo-loop*. (A) Stopped-flow analysis of the rates of Zn-cofactor binding to the pentacoordinate site. Lines drawn are fits with equation 5 with kon values of 1.8, 3.7, and 6.6 mM^-1^s^-1^ and koff values of 0.137, 0.112, and 0.062 s^-1^ for Zn-PPIX, ZnPcS, and ZnTPPS4 respectively. (B) Spectroscopic analysis of diheme tricofactor complexes. Oxidized (black lines) and reduced (red lines) absorbance (solid lines) and normalized fluorescence emission spectra in both oxidation states (black and red dashed lines) of diheme *photo-loop*:Zn-cofactor complexes. Fluorescence was measured using excitation wavelengths of 550 nm, 670 nm, and 570 nm, for ZnPPIX, ZnPcS, and ZnTPPS_4_ respectively. Upon heme oxidation, the fluorescence of these tricofactor complexes decreases 58%, 5%, and 29%.

**Table 2.**
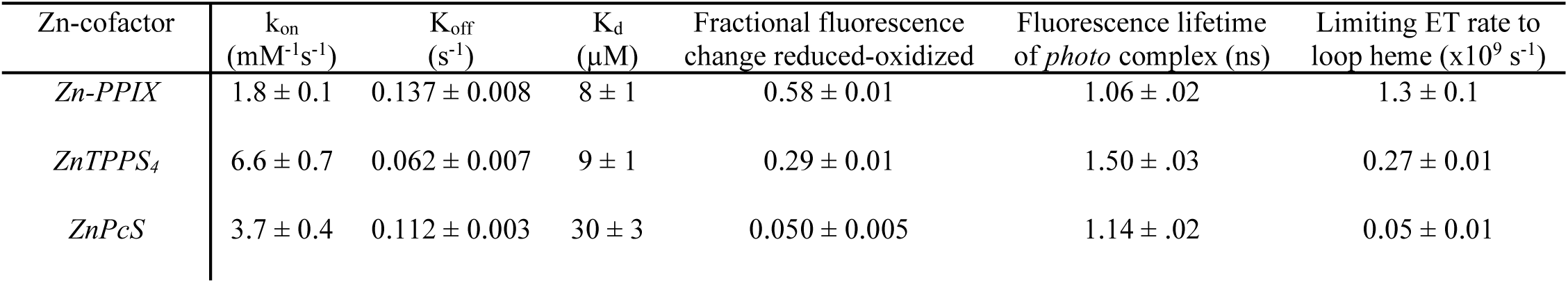
Thermodynamic and kinetic parameters for Zn-cofactors binding to diheme *photo-loop*.

| Zn-cofactor | k<sub>on</sub><br>(mM<sup>-1</sup>s<sup>-1</sup>) | K<sub>off</sub><br>(s<sup>-1</sup>) | K<sub>d</sub><br>(μM) | Fractional fluorescence<br>change reduced-oxidized | Fluorescence lifetime<br>of *photo* complex (ns) | Limiting ET rate to<br>loop heme (x10<sup>9</sup> s<sup>-1</sup>) |
|---|---|---|---|---|---|---|
| *Zn-PPIX* | 1.8 ± 0.1 | 0.137 ± 0.008 | 8 ± 1 | 0.58 ± 0.01 | 1.06 ± .02 | 1.3 ± 0.1 |
| *ZnTPPS<sub>4</sub>* | 6.6 ± 0.7 | 0.062 ± 0.007 | 9 ± 1 | 0.29 ± 0.01 | 1.50 ± .03 | 0.27 ± 0.01 |
| *ZnPcS* | 3.7 ± 0.4 | 0.112 ± 0.003 | 30 ± 3 | 0.050 ± 0.005 | 1.14 ± .02 | 0.05 ± 0.01 |

### Creating protein complexes with three different cofactors and quantifying intercofactor FRET

After the addition of a single heme, *photo-loop* binds two equivalents of any of the Zn-containing cofactors. Sequential addition experiments suggest that after the single heme is added, the first Zn cofactor binds to the internal binding site and the second binds to the loop binding site. One equivalent of Zn(II)-protoporphyrin IX (ZnPPIX) followed by the addition of a second molecule of heme does not interfere with a second heme binding, but two equivalents significantly increase the apparent heme dissociation constant. This means that the first ZnPPIX binds to the helical site and the second to the loop binding site. This enables us to create heterotricofactor complexes in which we can target cofactors to particular binding sites. First, a heme cofactor is added and binds to the internal bis-histidine binding site; next, a Zn cofactor is added and binds to the other internal binding site; last, another Zn-cofactor is added and binds to the loop histidines.

Figure 5 depicts the stepwise assembly of two different heterotricofactor complexes: one in which heme, Zn-PPIX and Zn(II)-phthalocyanine monosulphonate (ZnPcS) were added sequentially and one in which heme, Zn(II)-tetraphenzylporphyrin tetrasulfonate (ZnTPPS_4_) and ZnPcS were added sequentially. In each case, the addition of the second cofactor at the internal pentacoordinate binding site (red lines) results in minimal changes to the absorption spectrum, and thus the local environment, of the first cofactor as demonstrated by difference spectroscopy. Likewise, the addition of the third cofactor (blue lines) at the loop binding site has little effect on the spectra of the two bound cofactors.

**Figure 5.**
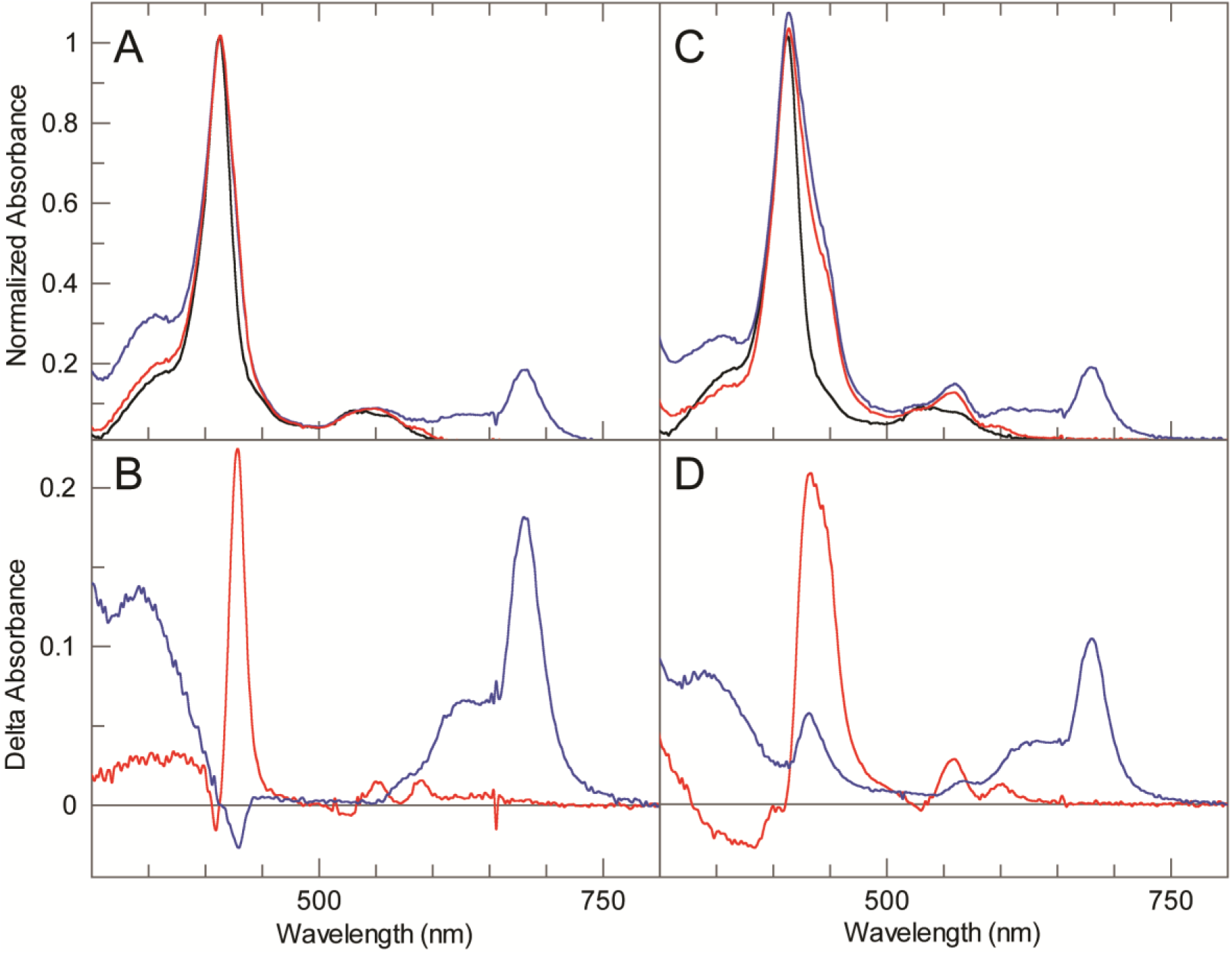
Stepwise assembly of two heterotricofactor complexes. (A) Absorption spectra of the photo-heme complex (black line), the heterobicomplex formed by the addition of one equivalent of ZnPPIX (red line), and the heterotricomplex formed by the further addition of ZnPcS (blue line). (B) Absorption difference spectra of the spectra from panel A depicting the sequential addition of ZnPPIX (red line) and and ZnPcS (blue line). (C) Absorption spectra of the photo-heme complex (black line), the heterobicomplex formed by the addition of one equivalent of ZnTPPS4 (red line), and the heterotricomplex formed by the further addition of ZnPcS (blue line). (D) Absorption difference spectra of the spectra from panel C depicting the sequential addition of ZnTPPS4 (red line) and and ZnPcS (blue line).

Fluorescence analyses of this stepwise addition process (Figure 6) demonstrate that both heterotricofactor complexes are functional light-activated energy-transfer proteins. Both dyads display fluorescence emission from the single Zn cofactor they contain. After the addition of the third photoactive cofactor, the original emission is reduced in intensity and longer-wavelength emission bands are observed from the third cofactor, although excitation wavelengths were used that do not directly excite the third cofactor. Quantitation of the emission intensity indicates 0.39 and 0.14 fractional energy transfer in the heme:Zn-PPIX:ZnPcS and heme: ZnTPPS_4_:ZnPcS complexes, respectively.

**Figure 6.**
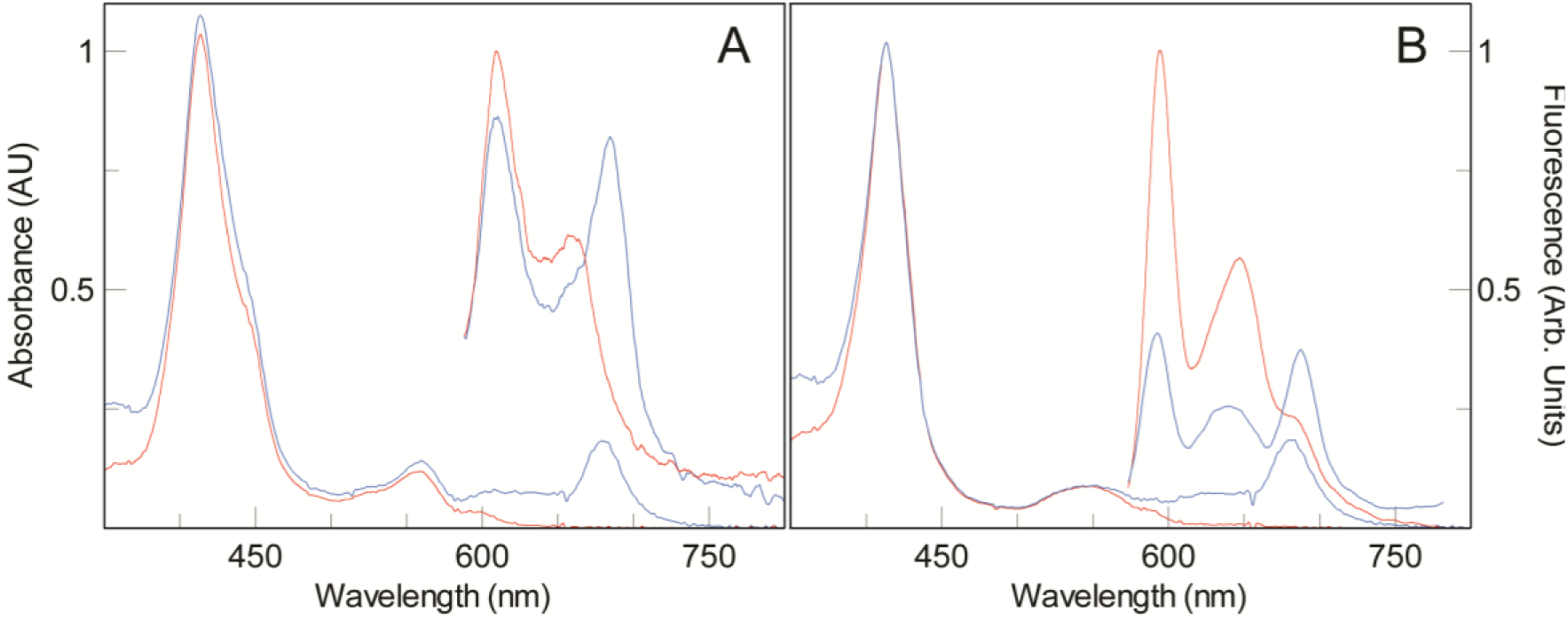
Excited state energy transfer between bound cofactors within two different heterotricofactor complexes. (A) Absorbance and fluorescence of 3 μM photo-loop protein bound with single equivalents of heme and ZnPPIX (red lines) and the same complex after ZnPcS is added (blue lines). The decrease in ZnPPIX fluorescence (595 and 647 nm) and increase in ZnPcS (688 nm) shows energy transfer within the protein sample when excited at 410 nm, an excitation wavelength which does not directly excite ZnPcS. (B) Absorbance and fluorescence of 3 μM photo-loop bound with one equivalent heme and ZnTPPS4 (red lines) and the same complex after ZnPcS is added (blue lines). The decrease in ZnTPPS4 fluorescence (609 nm) and increase in ZnPcS (685 nm) shows energy transfer sample when excited at 430 nm, an excitation wavelength which does not directly excite ZnPcS.

## Conclusion

Natural cofactor binding sites are known to occur predominantly in loop regions of proteins (21), yet most designed cofactor-binding proteins rely on helices or β-sheets for ligand coordination. Here we show that adding just two histidines to adjacent interhelical loops creates an additional site that binds ferric heme with an equilibrium dissociation constant of 90 nM. The combination of nanomolar affinity and minimal sequence modification establishes a simple route to expanding the cofactor-binding capacity of designed proteins. This contrasts sharply with our earlier bioinformatic analysis of helical heme- and porphyrin-binding motifs, which require five to seven precisely positioned residues to form a single complementary pocket (29). The effectiveness of this minimalist loop-based site helps explain why cofactors so often reside in loops in natural proteins: only limited packing complementarity is needed, meaning that just a few mutations can introduce a functional cofactor site, after which additional mutations can tune affinity, reactivity, and specificity.

A persistent barrier in engineering complex functional proteins is achieving controlled assembly of multiple cofactors within one domain. Here we have created and characterized several artificial, self-assembling heterotricofactor protein complexes using a design strategy so simple that it requires no computational optimization. These loop-embedded cofactor sites will allow us—and the broader field—to build more intricate functional protein assemblies than currently possible, enabling new approaches to modeling and understanding natural energy- and electron-transfer processes.

## Experimental

### Chemicals

Heme (Fe(III)-protoporphyrin IX) was purchased from Fluka (Buchs, Switzerland). Zn(II)-phthalocyanine (ZnPcS)^1^ was synthesized and purified as before (40). Zn(II)-tetraphenyl porphyrin tetrasulfonate (ZnTPPS_4_) and Zn(II)-protoporphyrin IX (ZnPPIX) were purchased from Frontier Scientific (Logan, UT). 1.0 mg/mL stock solutions of cofactors were freshly prepared in dimethyl DMSO and used within four hours. Bovine catalase was purchased from Calbiochem (San Diego, CA) and *Aspergillus niger* glucose oxidase was purchased from Amresco (Solon, OH). Apomyglobin (apoMG) was prepared from horse heart myoglobin using the method of Yonetani (41). Molecular nitrogen (N_2_) (99.99%) gas was purchased from T.W. Smith (Brooklyn, NY) and scrubbed of residual O_2_ by passage through two bubblers filled with a reduced vanadium sulfate solution followed by another filled with water (42). PD-10 desalting columns were from GE Healthcare (Port Washington, NY). All other solvents and reagents were from either VWR or Sigma.

### Molecular Biology

Genes encoding Photo and Photo-loop (see Figure 1) were commercially synthesized (Biomatik Corp., Cambridge, ON) with an N-terminal tobacco etch virus (TEV) recognition site and inserted between the BamHI and XhoI restriction sites of pET32a(+) (Merck Co., Darmstadt, Germany) creating expression plasmids which express His_6_-terminated thioredoxin fusion proteins with a TEV recognition site immediately before the desired protein. Further modifications were created by site-directed mutagenesis using the Agilent (Santa Clara, CA) QuikChange Lightning kit using primers purchased from Integrated DNA Technologies (Coralville, IA).

### Protein Structure Prediction

Protein structures for apo-photoloop and photoloop bound to two or three iron heme molecules were predicted using AlphaFold3 (34). The protein sequence and the number of heme molecules were provided as input, and the modeling was performed using the default settings, with the seed set to 123 for reproducibility. For each model, the placement of heme cofactors relative to the designed histidine ligands was inspected, and local model confidence was evaluated using the reported pLDDT values. Cofactor-associated pLDDT values were calculated as the average pLDDT for the heme atoms over five AlphaFold runs for each ligand state.

### Protein expression and purification

Artificial cofactor-binding proteins were grown and purified via native hexa-histidine tag purification as before (18). Briefly, 2L cultures of BL21(DE3) *Escherichia coli* containing the appropriate plasmid were grown in supplemented TPP media (43) at 298 K to an OD_600_ of 1.0 and then induced with 0.5 mM isopropyl-β-thiogalactopyranoside for 5 hours at room temperature. Cells were collected by centrifugation, broken open using a French press, and purified on a Ni-nitrilotriacetic acid column (Qiagen, Inc.) according to manufacturer’s instructions. The fusion protein was dialyzed into 50 mM Tris-HCl, 1 mM dithiothreitol, pH 8.0 and then cleaved overnight with His_6_-tagged TEV protease (44). The reaction mixture was filtered through Ni-nitrilotriacetic acid resin to remove the thioredoxin and protease, and the purified protein concentrated by centrifugation in Centricon YM-3 centrifugal filter units (EMD-Millipore, Inc., Billerica, MA) when necessary. All stages of purification were monitored by SDS-PAGE. Final purity was confirmed using analytical reversed-phase C18 high-pressure liquid chromatography. Pure proteins were dialyzed into 250 mM boric acid, 100 mM KCl, pH 9.0 unless otherwise specified.

The gene for *E. coli.* ferredoxin-NADP^+^ reductase in expression plasmid pET28a was a kind gift from Professor Mark Hargrove, Iowa State University Department of Biochemistry, and was purified by standard native nickel column purification procedures from BL21 *E. coli* using the literature procedure (45). Ferredoxin-NADP^+^ reductase activity was assayed using the cytochrome *c* reductase assay (46).

### General biochemistry

Absorbance spectra were collected with a Hewlett-Packard (New York, NY) 8452A Diode array spectrophotometer running the Olis (Bogart, GA) SpectralWorks software and equipped with a Quantum Northwest (Liberty Lake, WA) Peltier temperature controller. All experiments were performed at 298K in 250 mM boric acid, 100 mM KCl, pH 9.0 unless otherwise specified. Each binding and reduction potential experiment was performed at least three times, and reported values are the average and errors are standard deviations from the mean.

Fluorescence emission spectra were collected on an Olis DM45 dual monochromator fluorescence spectrometer equipped with a Quantum Northwest temperature controller maintained at 298K. Cofactor fluorescence intensities were recorded with 0.6 nm and 1.24 nm slits width for excitation and emission respectively. Excitation wavelengths of 426 nm, 600 nm and 630 nm were used for ZnPPIX, ZnTTPS_4_ and ZnPcS, respectively.

Cofactor-protein complexes were prepared as before (16) by consecutive additions of 0.2 equivalents of cofactor in DMSO up to the desired fractional cofactor loading with at least ten minutes between additions. After cofactor loading, proteins were then isolated from any unbound cofactor using PD10 desalting columns. Multicofactor proteins were prepared by sequential addition of cofactors to 2.5 mL solution in borate buffer as described above with desalting between cofactor additions. For three-cofactor assemblies, the third cofactor bound was always the primary donor cofactor, loading was limited to 0.8 equivalents, and spectral analyses were performed directly without desalting.

When anaerobicity was needed, O_2_ removal was performed using the glucose oxidase/catalase enzymatic system described by Benesch *et al.* using 2% glucose with 25 units/mL glucose oxidase and 250 units/mL catalase (47; 48) in the cuvette. During experiments with ferrous heme cofactors, proteins were reduced using 0.20 mM NADPH and 0.25 µM ferredoxin-NADP reductase.

### Heme binding affinity measurements

Protein solutions were 5-10 µM in 250 mM borate buffer, pH 9.0. Experiments were performed in 1 cm path-length screw-top cuvettes. Hemin solutions were approximately 1mg/mL in DMSO and exact concentrations were determined using the pyridine hemochrome method of Trumpower (49). Successive 0.1-0.2-equivalent additions of cofactor were followed by a 20-minute equilibration period at 20°C. Visible spectra were collected after each equilibration and binding was monitored using the bound heme Soret peak at 412 nm for oxidized heme and 426 nm for reduced heme. Binding affinities of reduced heme were determined in anaerobic cuvettes under reducing conditions as performed previously (16). K_d_ values were obtained from plots of the Soret band absorbance measured at 426 nm vs. the concentration of hemin added and fit with the tight binding equation:

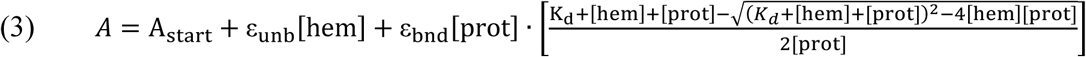

Where ε_unb_ is the molar absorption coefficient of unbound hemin at that wavelength, ε_bnd_ is the additional absorbance of bound hemin at that wavelength, [hem] is the hemin concentration, [prot] is the protein concentration and K_d_ is the dissociation constant for the reduced hemin.

### Zn-cofactor binding

For ZnPPIX, ZnTPPS_4_ and ZnPcS binding, protein samples were first bound to a single equivalent of heme as above. After addition of a full equivalent of heme, these samples were then passed over a PD-10 desalting column (GE Healthcare, Piscataway, NJ) to remove excess DMSO. Protein-heme complex concentrations were calculated using the bound heme Soret peak at 412 nm.

Quantitative ZnPcS and ZnTPPS_4_ binding titrations were performed by adding freshly prepared cofactor solution in dimethylformamide to anaerobic 1 mL aliquots of protein-heme complex in 0.1 equivalent quantities at 20-minute intervals. Binding was monitored through spectral changes in the phthalocyanine and porphyrin Q bands (630-700 nm)(50–52) and the data were fit with equation 1. For fluorescence spectroscopy measurements protein-heme-Zn cofactor complexes were similarly prepared and purified using a PD-10 column as above.

Anaerobic Zn(II)-PPIX binding was quantified via the differential fluorescence emission at 598 nm between bound and free cofactor. Emission data were fit with equation 4:

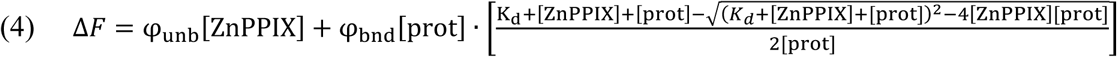

Where φ_unb_ is the fluorescence emission intensity of unbound ZnPPIX and φ_bnd_ is the fluorescence emission intensity of bound ZnPPIX. The response of the fluorimeter was externally calibrated before each Zn(II)-PPIX titration using the fluorescence of 4-(dicyanomethylene)-2-methyl-6-(4-dimethylaminostyryl)-4*H*-pyran at identical excitation and emission wavelengths.(53) ZnPPIX-protein complex fluorescence data are normalized to the molar emission intensity of the external standard.

### Stopped-flow analysis of ligand binding rates

Binding kinetics of cofactors were followed spectroscopically in rapid stopped-flow mixing experiments at 20° C using a Biologic (Lyon, France) SFM 400 stopped flow mixer equipped with a custom-built Olis RSM 1000 spectrometer for multiwavelength absorption detection. Heme binding rate determination was performed using the Olis RSM 1000 with a 0.6 cm cuvette path length and 0.6 nm slit width, sampling the Soret band spectral absorption region centered at 420 nm. Heme working solutions were prepared by dissolving stock solutions prepared in 25 mM NaOH into buffer immediately before data acquisition. Zinc cofactor binding rate determination was performed using fluorescence detection in a 1.0 x 0.6 cm path length cuvette with 3.16 nm excitation and emission slit widths with excitation at 408 nm and a 500 nm high-pass optical filter. Stock solution of fluorescent cofactors in DMSO were dissolved immediately before use. Final concentrations after mixing were 1 µM protein and 2-16 µM cofactor. Binding data were fit with single exponentials and the combined data were fit with:

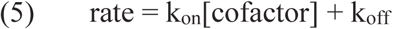

where k_on_ and k_off_ are the association and dissociation rate constants, respectively.

### Stopped flow analysis of heme dissociation

Dissociation rates for the bis-histidine–ligated hemes in both heme-*photo* and diheme *photo-loop* constructs were determined by trapping dissociated hemes with apoMG (54; 36). Final protein concentrations after mixing were 6 μM heme-photo or photo-loop and 26 or 52 μM apoMb. For the photo-loop construct, the loop-binding site was only half-loaded to ensure complete binding of the loop heme site. Heme off-rates were obtained by monitoring the increase of the bis-histidine Soret absorbance at 426 nm, which increases upon formation of the five-coordinate deoxy-myoglobin–heme complex. Data for the heme*-photo* construct were fit to a single exponential. Data for the *photo-loop* construct were fit to:

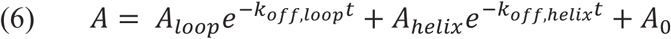

where A_loop_ is the absorbance at 412 nm of the loop bound heme, k_off,loop_ is the dissociation rate of the loop heme, A_helix_ and k_off,helix_ are the absorbance and dissociation rate of the helical heme.

### Bound heme reduction potential determination

Potentiometric oxidation/reduction titrations were performed in combination with optical analysis as described previously (55). Reported reduction potentials are referenced to a standard hydrogen electrode. All redox titrations were performed anaerobically using *μ*L additions of freshly prepared sodium dithionite to adjust the solution potential to more negative values and potassium ferricyanide to adjust it to more positive values. Redox titrations were analyzed by monitoring the Q absorbance band at 528 nm as the heme protein was reduced or oxidized. The data were analyzed with the Nernst equation using an *n*-value of 1.0:

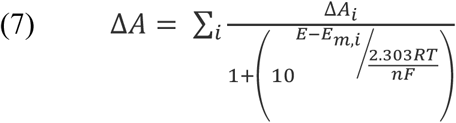

where ΔA_i_ is the change in absorbance at either 560 or 528 nm for each heme during the titration, E is the solution potential, E_m,i_ is the reduction midpoint potential for each heme, *n* is the number of electrons, F is Faraday’s constant, and *i* is the cofactor index.

### Zn-heme fluorescence lifetime

Time-resolved photoluminescence measurements of Zn-heme samples were acquired using a supercontinuum fiber laser (NKT Photonics, Super K) operating at 5 MHz as the excitation source. Samples were excited at 426 nm and a Hamamatsu C10910−04 streak camera was used to collect the spectra. Appropriate filters were used to eliminate scattered excitation light from the PL measurements. The lifetime was calculated from single exponential fits of the decay and the reported error is the error of the fit.

### Light-activated electron transfer

Electron transfer rates from photoexcited Zn-heme complexes containing acceptor residues or cofactors were calculated using the Stern-Vollmer relationship:(22)

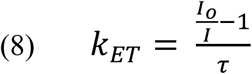

Where I_o_ and τ are the steady-state fluorescence and fluorescence lifetime of the *photo*:Zn-heme complex, respectively, and I is the steady-state fluorescence of the complex in question. *Energy transfer measurements.* Intercofactor resonance energy transfer measurements were performed by exciting ZnPPIX and ZnTTPS_4_ donor assemblies at 424 nm and 430 nm respectively and observing the acceptor cofactor ZnPcS fluorescence at 685 nm. Entry and exit slits for both monochromators were fixed at 1.24 nm. Freshly prepared protein was prepared by sequential addition of heme followed by ZnPPIX or ZnTTPS_4_ as described previously. Fluorescence measurements were taken before ZnPcS addition and after 0.8 equivalents were added to the cuvette with dyad protein. Fluorescence emission was normalized to greatest intensity. Excitation scans (400-625 nm) were performed at the same time on the same samples.

## Supporting information

Supplemental figures and derivation for Mutter et al

## Acknowledgements

The authors would like to thank Peter Tipton, of the University of Missouri Department of Biochemistry for helpful discussions regarding binding kinetics. RLK gratefully acknowledges support from the NSF (MCB-2025200). ACM gratefully acknowledges support from the Center for Exploitation of Nanostructures in Sensor and Energy Systems (CENSES) under NSF Cooperative Agreement Award Number 0833180. AU was the recipient of a fellowship award from the U.S. Department of Education Graduate Assistance in Areas of National Need (GAANN) Program in Biochemistry, Biophysics, and Biodesign at The City College of New York (PA200A150068). This work was authored in part by the National Laboratory of the Rockies for the U.S. Department of Energy (DOE), operated under Contract No. DE-AC36-08GO28308. Funding provided by U.S. Department of Energy Office of Basic Energy Sciences, Division of Chemical Sciences, Geosciences, and Biosciences, Physical Biosciences Program. The views expressed in the article do not necessarily represent the views of the DOE or the U.S. Government. The U.S. Government retains and the publisher, by accepting the article for publication, acknowledges that the U.S. Government retains a nonexclusive, paid-up, irrevocable, worldwide license to publish or reproduce the published form of this work, or allow others to do so, for U.S. Government purposes.

