## Supplemental figures and derivation for Mutter et al for "Loop-Directed Cofactor Binding Enables Programmable Multicofactor Protein Design"

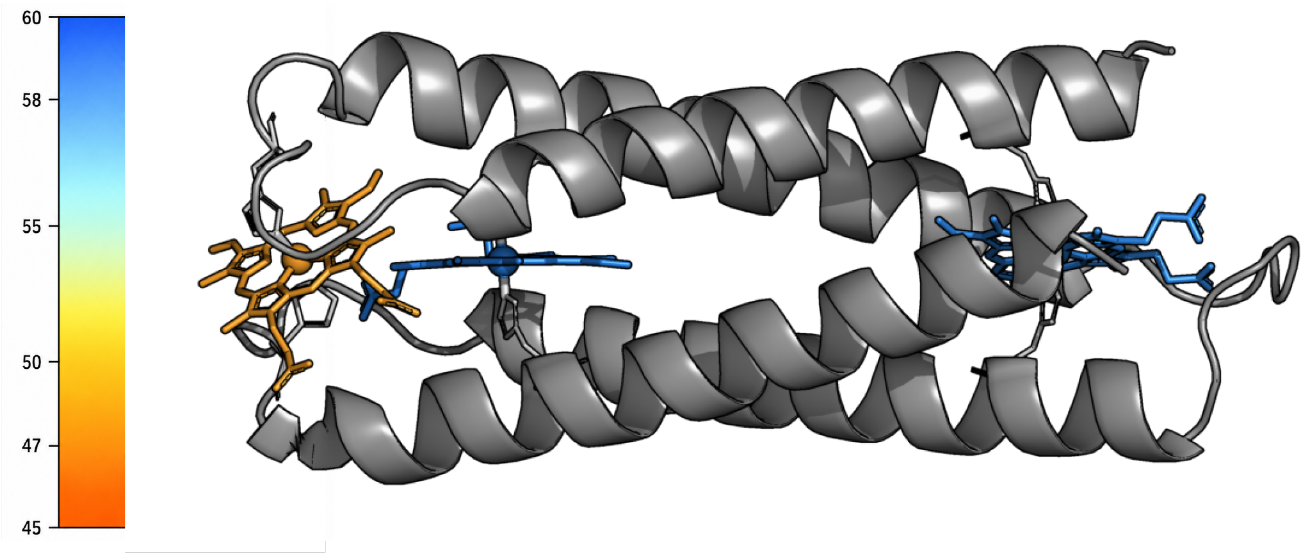


**Supplementary Figure S1.** AlphaFold3 model of predicted heme cofactor binding (Right) Representative three-heme model showing the protein in grey ribbon with explicit histidines and the three modeled heme cofactors as sticks. Heme cofactors are colored by across-model mean ligand pLDDT using an intentionally compressed color scale (left) to emphasize relative confidence differences among the three sites. The open end heme and the Zn-heme binding sites have much higher confidences (58 ± 1), whereas the loop binding site has a lower mean ligand pLDDT (47 ± 2). The lower mean pLDDT of the loop-bound heme indicates greater uncertainty in its predicted placement relative to the buried cofactors; these scores describe prediction confidence in a binary-patterned scaffold that was developed independently of AlphaFold and thus have lower than the confidences seen in hologues or mutants of known structures.

**Supplemental Figure S2.** Calculation of distal histidine on- and off-rates. Constants in red are experimentally determined values.

$$K_d = \frac{k_{off}}{k_{on}}\left(\frac{k_{-His}}{k_{+His}+k_{-His}}\right)$$

$$\downarrow$$

$$\frac{K_d k_{on}}{k_{off}} = \left(\frac{k_{-His}}{k_{+His}+k_{-His}}\right)$$

$$\downarrow$$

$$\frac{K_d k_{on}}{k_{off}}(k_{+His} + k_{-His}) = k_{-His}$$

$$\downarrow$$

$$\frac{K_d k_{on}}{k_{off}} k_{+His} = k_{-His}\left(1 - \frac{K_d k_{on}}{k_{off}}\right)$$

$$\downarrow$$

$$k_{+His} = k_{-His}\left(\frac{k_{off}}{K_d k_{on}} - 1\right)$$

and

$$k_{off,app} = \frac{k_{off} k_{-His}}{k_{off}+k_{+His}}$$

$$\downarrow$$

$$\downarrow$$

$$\downarrow$$

$$k_{off,app} = \frac{k_{off} k_{-His}}{k_{off}+k_{-His}\left(\frac{k_{off}}{K_d k_{on}}-1\right)}$$

$$\downarrow$$

$$k_{off,app}\left(k_{off} + k_{-His}\left(\frac{k_{off}}{K_d k_{on}} - 1\right)\right) = k_{off} k_{-His}$$

$$\downarrow$$

$$k_{off,app} k_{off} + k_{-His} k_{off,app}\left(\frac{k_{off}}{K_d k_{on}} - 1\right) = k_{off} k_{-His}$$

$$\downarrow$$

$$k_{off,app} k_{off} = k_{off} k_{-His} - k_{-His} k_{off,app}\left(\frac{k_{off}}{K_d k_{on}} - 1\right)$$

$$\downarrow$$

$$k_{off,app} k_{off} = k_{-His}\left[k_{off} - k_{off,app}\left(\frac{k_{off}}{K_d k_{on}} - 1\right)\right]$$

$$\downarrow$$

$$k_{-His} = \frac{k_{off,app} k_{off}}{k_{off} - k_{off,app}\left(\frac{k_{off}}{K_d k_{on}} - 1\right)}$$

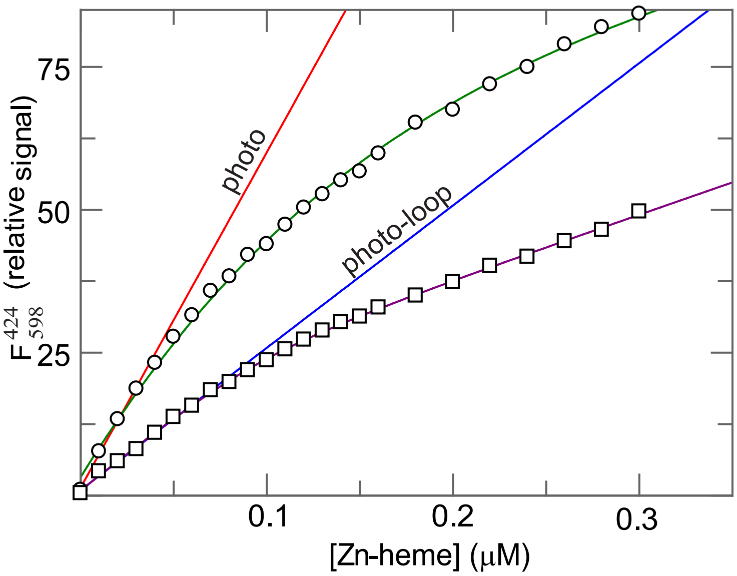


**Supplementary Figure S3.** Fluorescence titration of ZnPPIX binding to 100 nM solutions of monoheme photo and diheme photo-loop. ZnPPIX fluorescence was monitored at 598 nm following excitation at 424 nm during titration into heme-loaded photo (circles) and diheme photo-loop (squares). Curved lines are fits with the tight-binding equation (Eq. 4) with dissociation constants of 10 ± 2 nM for photo and 8 ± 5 for photo-loop. Straight lines are a linear fit of the first 0.2 equivalents with slopes of 590 ± 20 and 250 ± 30 $\mu M^{-1}$, respectively. The ratio of slopes in combination with Eq. 8 predicts a rate of (1.3 ± 0.2) $x10^9$ s-1, within error of the lifetime calculated using the data in Figure 4B.
